# A Convolutional Neural Network and R-Shiny App for Automated Identification and Classification of Animal Sounds

**DOI:** 10.1101/2020.07.15.204685

**Authors:** Zachary J. Ruff, Damon B. Lesmeister, Cara L. Appel, Christopher M. Sullivan

## Abstract

The use of passive acoustic monitoring in wildlife ecology has increased dramatically in recent years as researchers take advantage of improvements in automated recording units and associated technologies. These technologies have allowed researchers to collect large quantities of acoustic data which must then be processed to extract meaningful information, e.g. target species detections. A persistent issue in acoustic monitoring is the challenge of processing these data most efficiently to automate the detection of species of interest, and deep learning has emerged as a powerful approach to achieve these objectives. Here we report on the development and use of a deep convolutional neural network for the automated detection of 14 forest-adapted species by classifying spectrogram images generated from short audio clips. The neural network has improved performance compared to models previously developed for some of the target classes. Our neural network performed well for most species and at least satisfactory for others. To improve portability and usability by field biologists, we developed a graphical interface for the neural network that can be run through RStudio using the Shiny package, creating a highly portable solution to efficiently process audio data closer to the point of collection and with minimal delays using consumer-grade computers.

## Introduction

Artificial intelligence (AI) technologies are increasingly being applied to issues in ecological research and conservation. In the field of wildlife ecology, the use of AI in combination with recent advances in survey techniques has enabled researchers to collect data on species occurrences at much broader spatial and temporal scales than were previously possible. Passive monitoring methods such as camera traps and bioacoustics have greatly improved the capacity of researchers to survey for wildlife, but the resulting large datasets require substantial processing to extract useful information. The task of quickly and accurately locating signals of interest (e.g., target species detections) within large audio or photo datasets remains a persistent issue. Automated or semi-automated approaches can greatly reduce the amount of time and effort required to extract and classify detections of target animals (Norouzzadeh et al. 2017, Willi et al. 2018).

Recent efforts have successfully employed image recognition algorithms to describe, count, and identify animals of interest in passively collected data (Weinstein 2017). Deep convolutional neural networks (CNN), a deep learning approach commonly used for image recognition, have proven particularly useful in analysis of ecological data (Brodrick et al. 2019, Ruff et al. 2020). Several researchers have used CNNs to identify and classify animal species in images from remote camera traps with up to 98% accuracy (Gomez Villa et al. 2017, Norouzzadeh et al. 2017, Tabak et al. 2019).

Computer vision techniques can also be used to extract detections of vocalizing or otherwise audible species from acoustic recordings. By converting sound files into spectrograms—a visual representation of the energy present over a range of acoustic frequencies as a function of time—data recorded by acoustic recording devices may then be classified using the same algorithms that make image recognition from camera trap photos possible. Recordings of avian vocalizations have been analyzed using a variety of automated recognizer algorithms (e.g., Sebastián-González et al. 2015, Knight et al. 2017, Venier et al. 2017, Ovaskainen et al. 2018, Stowell et al. 2019), including CNNs specifically (Salamon et al. 2016, Salamon and Bello 2017, Ruff et al. 2020, LeBien et al. 2020).

Ruff et al. (2020) developed a CNN that identified vocalizations of six owl species in large sets of unprocessed audio recordings generated by autonomous recording units (ARUs) for the purpose of passive acoustic monitoring of wildlife. In this paper we report on advancements to the CNN that was developed by Ruff et al. (2020), including the incorporation of 10 additional target classes and improved performance on the six owl species classes. We used the CNN to process audio data recorded in forested landscapes of Oregon and Washington, USA. Human verification of apparent detections was an integral part of the process; however, model classification performance has improved greatly, and our verification methods were refined to the point where only cursory human effort was necessary to confirm the presence of target species at a site. This process enabled us to quickly derive weekly encounter histories from ARU data, to be used for occupancy modeling and other analyses.

As detailed in Ruff et al. (2020), the project derived from ongoing monitoring efforts for the northern spotted owl (*Strix occidentalis caurina*), which is an important conservation and management species in the Pacific Northwest, USA, and is facing declining populations and various threats to persistence (Lesmeister et al. 2018, Jenkins et al. 2019, Wiens et al. 2019). Passive acoustic monitoring has been demonstrated to be effective at detecting rare species such as the northern spotted owl (Duchac et al. 2020), prompting the deployment of large-scale acoustic monitoring for this and other species (Lesmeister et al. 2019).

Techniques for automatic detection of target species often require substantial computing power, and data processing is often conducted after the end of the field season to take advantage of high-performance computers, but this approach creates significant delays between data collection and analysis. A secondary objective to our CNN training was to develop a tool that would allow biologists and managers in the field to process data in a distributed fashion, close to the point of collection and with minimal delays in the output of detections. To this end, we packaged the CNN as a desktop application with a Shiny graphical interface which can be run through RStudio (RStudio Team 2020), creating a highly portable solution that allows efficient processing of audio data on consumer-grade desktop computers.

The Ruff et al. (2020) CNN had seven target classes, including six species of owls: barred owl *Strix varia*, great horned owl *Bubo virginianus*, northern pygmy-owl *Glaucidium gnoma*, northern saw-whet owl *Aegolius acadicus*, northern spotted owl, and western screech-owl *Megascops kennicottii*, as well as a catch-all “Noise” class to encompass any image not classifiable as a target species. For our CNN we included those seven classes but expanded the class list to include eight additional target species: band-tailed pigeon *Patagioenas fasciata*, common raven *Corvus corax*, Douglas’ squirrel *Tamiasciurus douglasii*, mountain quail *Oreortyx pictus*, pileated woodpecker *Dryocopus pileatus*, red-breasted sapsucker *Sphyrapicus ruber*, Steller’s jay *Cyanocitta stelleri*, and Townsend’s chipmunk *Neotamias townsendii*.

Some of the target classes were included because the species in question were of ecological or management interest due to potential competitive or predatory interactions with other species. For example, Townsend’s chipmunks are important prey for many raptors and mammalian predators. Other classes were added because previous verification efforts indicated that they were likely to produce false-positive detections for existing target classes. Some target species fulfilled more than one of these criteria; band-tailed pigeon is a managed game species (Sanders 2015) that was extremely common at our study sites, and was a major source of false positive detections for great horned owl in the Ruff et al. (2020) study, and Townsend’s chipmunk calls are easily confused with northern saw-whet owl calls.

## Methods

### Training data

Our training dataset included 53,292 unique clips of vocalizations from 14 species (Table 1). For eight of these species, including band-tailed pigeon, great horned owl, mountain quail, northern pygmy-owl, northern saw-whet owl, northern spotted owl, red-breasted sapsucker, and Townsend’s chipmunk, the examples in the training set included only one highly-stereotyped call type or sound. For another four species, namely common raven, pileated woodpecker, Steller’s jay, and western screech-owl, the training set included multiple call types, but the call types for each species were lumped into one class because the component syllables were sufficiently similar that including closely related call types would likely not hinder identification. For the remaining two species, barred owl and Douglas’ squirrel, we included two call types but incorporated each call type as a separate class. Like Ruff et al. (2020) we included a catch-all “Noise” class for any image without one of our target species, which we reasoned would improve CNN performance for our target classes.

**Table 1.**
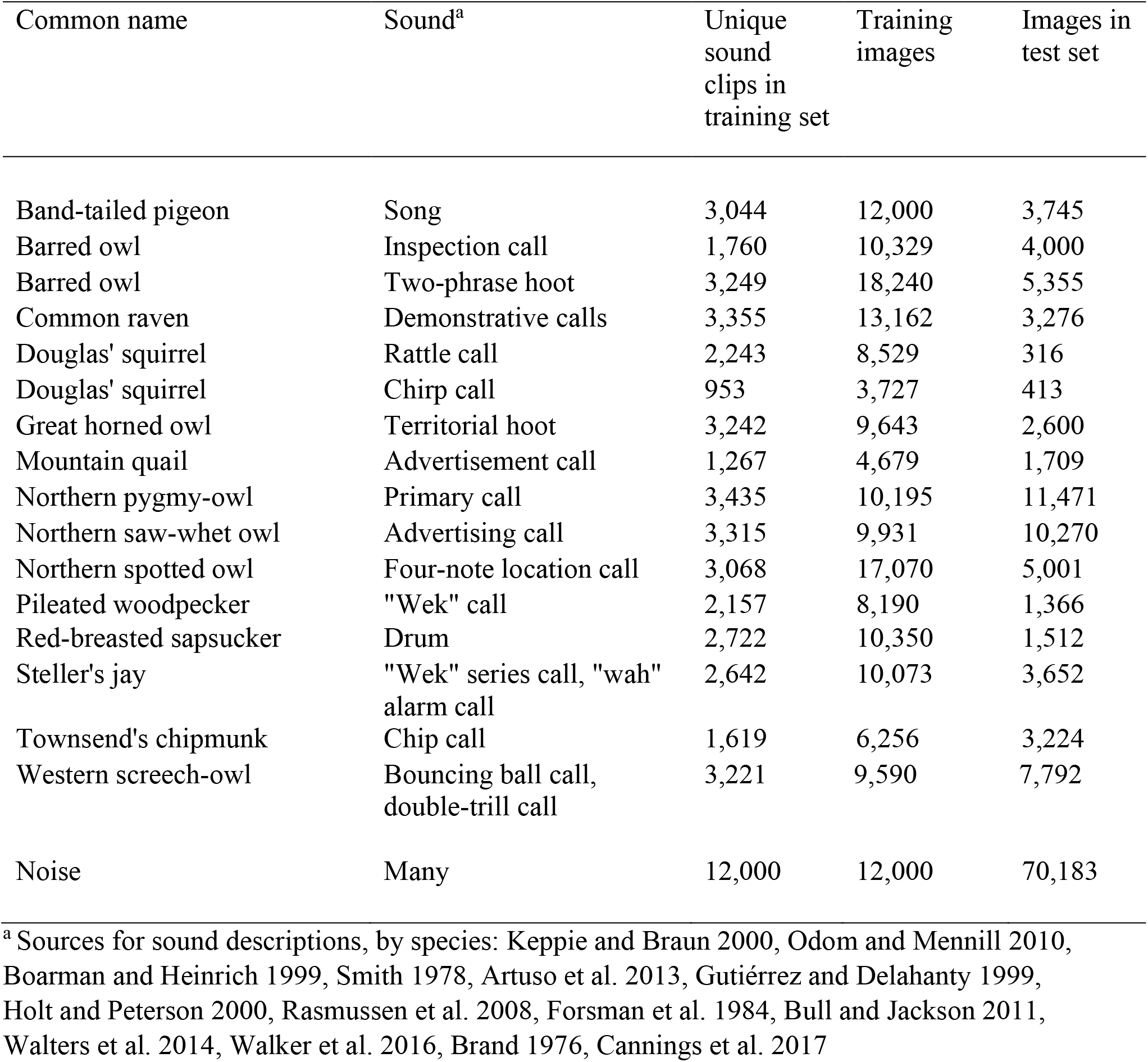
Target species and the characteristic sounds used to train the convolutional neural network and construct the test set. Each row denotes a separate class and corresponds to one node in the network’s output layer. We had a number of unique audio clips representing each class, from which we generated spectrogram images that we used to train the network. Each class was a specific call type or group of vocalizations with similar syllables. We generated three to six variant spectrograms with slightly different parameters for each clip to increase the volume of training data. During processing, long audio clips are segmented into spectrograms, each representing 12 seconds of audio. For each image the trained network outputs a vector of 17 class scores, each between zero and one, representing the strength of the match between the image and each of the target classes. The test set (*n* = 131,767) was comprised of images generated from examples of the same call types that were not used to generate spectrograms for the training or validation set. Some images in the test set contained calls from >1 target class.

Training images were generated from annotated records of calls found in data from a survey effort for northern spotted owls and barred owls (see Duchac et al. 2020). We reanalyzed those data and searched semi-manually for calls using the simple clustering feature of Kaleidoscope Pro software (version 5.0, Wildlife Acoustics). The passive acoustic monitoring occurred during Mar – July 2017 on 150 sites on three historical northern spotted owl demographic study areas (Coast Range, Klamath, Olympic Peninsula) in the Pacific Northwest, USA. See Forsman et al. (2011) for study area descriptions. For each class we included training examples from at least two and in most cases all three study areas, to ensure that the CNN’s internal representation of each class would be less influenced by systematic regional differences in vocalizations or background noise.

To augment the unique sound clips included in the training set we generated multiple variant spectrograms with randomized offset and dynamic range, producing three to six distinct images for each unique call, using the same procedure detailed in Ruff et al. (2020). Spectrograms (Fig. 1) consisted of grayscale images in portable network graphic (PNG) format with resolution of 500 × 129 pixels and a bit depth of eight. We generated these images using the spectrogram command in SoX (version 14.4, http://sox.sourceforge.net). After creating the images, we reviewed the training set to ensure that each image contained a visible signature of sounds corresponding to one of our target classes, but no other classes. We reserved a randomly selected 20% of these images for the validation set and used the other 80% of images for training the CNN. The training set included spectrograms used to train the Ruff et al. (2020) CNN, as well as many additional images for those original target species and the ten additional classes. Because many sounds that were previously considered “noise” corresponded to one of the new target classes, we reviewed images included in the previous training set again to remove any spectrograms that contained calls of multiple species. Following the generation of the variant spectrograms, the final training dataset comprised 173,964 images representing our 17 classes (Table 1).

**Figure 1.**
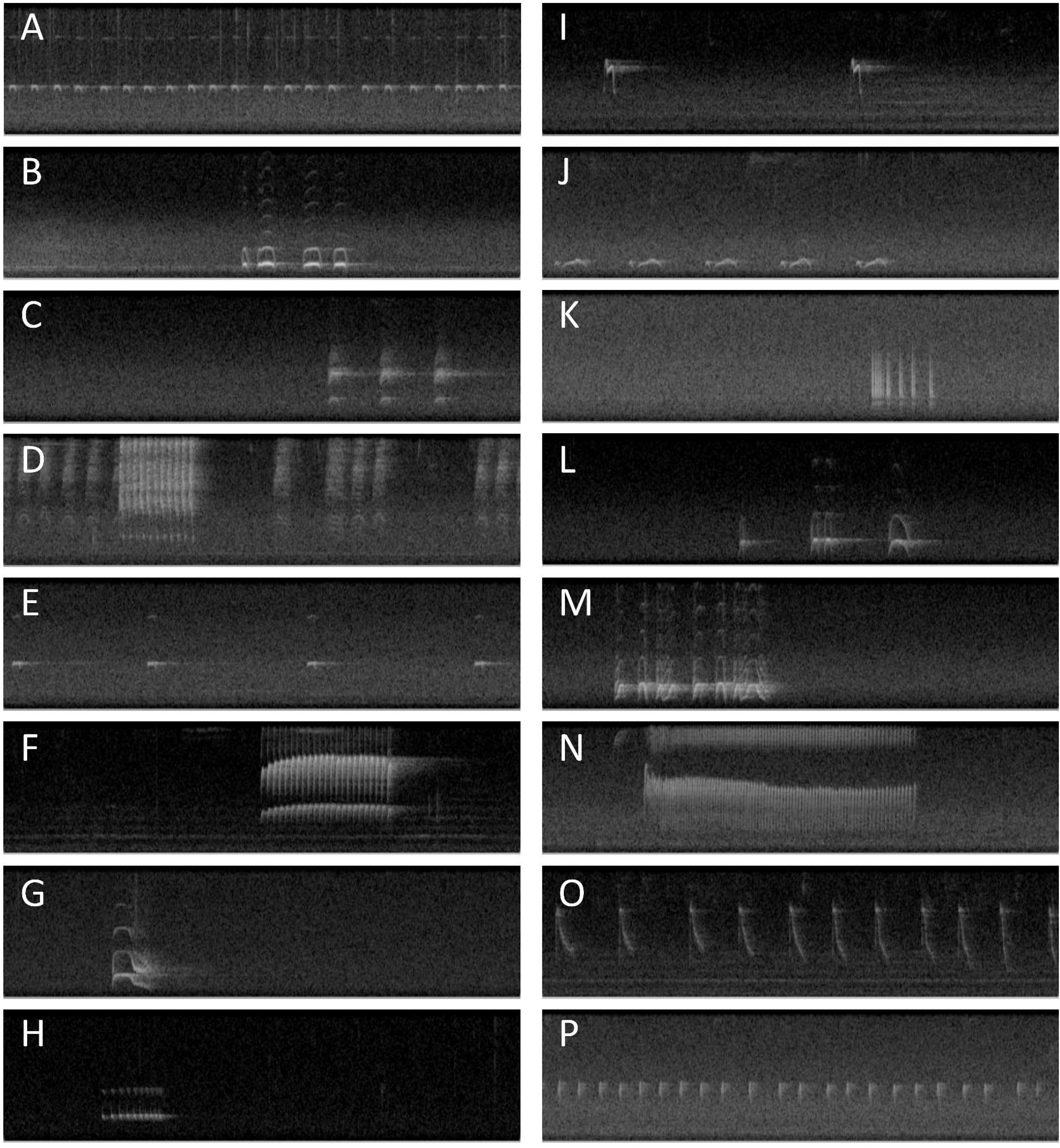
Examples of spectrograms representing each of 16 target classes. Spectrograms plot sound energy as a function of time (horizontal axis) and frequency (vertical axis); lighter colors represent higher energy intensity. These spectrograms represent 12 s of sound in the frequency range 0 – 3000 Hz. A. Northern saw-whet owl advertising call. B. Great horned owl territorial call. C. Common raven demonstrative calls. D. Steller’s jay “wek” series call and “wah” alarm call. E. Northern pygmy-owl primary call. F. Pileated woodpecker “wek” call. G. Barred owl inspection call. H. Western screech-owl “bouncing ball” call. I. Mountain quail advertisement call. J. Band-tailed pigeon song. K. Red-breasted sapsucker drum. L. Northern spotted owl four-note location call. M. Barred owl two-phrase hoot. N. Douglas’ squirrel rattle call. O. Douglas’ squirrel chirp call. P. Townsend’s chipmunk chip call. We also used a Noise class (not pictured) for images containing only background noise or sounds that did not correspond to any target class.

### Convolutional neural network training

We implemented the CNN model in Python (version 2.7, Python Foundation) using Keras (Chollet 2015), an open-source, machine learning-focused application programming interface to Google’s TensorFlow software library (Abadi et al. 2015). The CNN contained six trainable layers, including four convolutional layers and two fully connected layers. The first convolutional layer contained 32 5×5 filters, the second layer contained 32 3×3 filters, and the third and fourth layers each contained 64 3×3 filters. Each convolutional layer had a stride length of one and used Rectified Linear Unit (ReLU) activation. Each convolutional layer was followed by 2×2 max pooling and 20% dropout. The first fully connected layer contained 256 units using ReLU activation and L2 regularization and was followed by 50% dropout. The L2 regularization was the squared Euclidean norm of the weight matrix of the hidden layer, or the sum of all such squared norms, in the case of multiple hidden layers, and including the output layer. The use of regularization techniques such as L2 may have been redundant to other optimization techniques with adaptive learning rates (Loschilov and Hutter 2019). The second fully connected layer, which was the output layer of the model, contained 17 units using sigmoid activation. Hence, the output from the model was a 17-element vector of class scores, each of which was between zero and one. Sigmoid activation is not normalized and thus the class scores that comprise the output of this CNN do not sum to one for a given input. In contrast, the Ruff et al. (2020) model used softmax activation in the final layer, which constrained output scores to a sum of one. In a model with sigmoid activation, classes are not implied to be exhaustive of every sound on the landscape; this is a major difference between a model with softmax activation and one with sigmoid activation.

We trained the CNN for 100 epochs using a batch size of 128 images. We measured loss using the binary cross-entropy function and used the Adam optimization algorithm (Kingma and Ba 2015) with an initial learning rate of 0.001. To prevent overfitting we saved the model after epochs in which validation loss decreased. We also included a stepdown function to adjust the learning rate during training: if the validation loss did not decrease by at least 0.025 for five epochs, the learning rate was reduced by half. This was followed by a cooldown period of six epochs; hence, the learning rate could diminish at a maximum rate of once every ten epochs. We implemented the cooldown period based on the observation that improvements in model performance during training are stochastic and therefore it might take several epochs to realize the potential benefit of a given learning rate. We trained the CNN using an IBM POWER8 high-performance computer running the IBM OpenPOWER Linux-based OS with two Nvidia Tesla P100-SXM2-16GB general-purpose graphics processing units.

### Model testing

To evaluate the performance of the new model in a way that would require less human validation and would not be biased by the verification procedure itself, we compiled an independent test set of 131,767 images for which the correct labels were known and which had not been part of the training or validation set (Table 1). This collection of test set images contained examples of all 17 target classes, of which approximately half (*n* = 70,183) were in the Noise class. The test set included 4,082 images containing multiple target classes; of these, 4,047 contained two target classes, 34 contained three target classes, and one contained four target classes. Most of the calls included in the test set were located opportunistically during human verification of output from the Ruff et al. (2020) CNN. During this process we tagged images by identifying species by ear and by examination of the spectrogram. After assembling the test set we used the presented version of the CNN to classify the images, and we report performance metrics based on the class scores that it assigned to each image.

Here we report the same metrics that were reported by Ruff et al. (2020), namely precision, recall, and F1 score, and using the same definition of detection threshold, i.e. an image was considered a potential detection for a class if the class score for that image exceeded the threshold.. Precision is defined as True Positives / (True Positives + False Positives) and represents the proportion of apparent ‘hits’ that are real detections. Recall is defined as True Positives / (True Positives + False Negatives) and represents the proportion of real examples in the dataset that are detected and correctly labeled. F1 score is a balanced combination of precision and recall, intended to measure overall performance. The F-score can be weighted to emphasize either precision or recall; we report the unweighted version, calculated as (2 * Precision * Recall) / (Precision + Recall).

For easier comparison to other models and in keeping with recommendations by Knight et al. (2017) we present receiver operating characteristic (ROC) curves for each species, which plot the true positive rate (recall), calculated as [True Positives] / [True Positives + False Negatives] against the false positive rate, calculated as [False Positives] / [False Positives + True Negatives]. We calculated the Area Under the Curve (AUC) values for each species for the ROC curves, which was the probability that the model assigned a higher score to a randomly chosen positive example than to a randomly chosen negative example. We also plotted Precision-Recall (PR) curves for each species, which plot precision (True Positives / [True Positives + False Positives]) against recall (True Positives / [True Positives + False Negatives]) to illustrate a given model’s performance and the magnitude of the trade-off between these metrics across a range of detection thresholds. We calculated the AUC for the PR curves; these values had no intuitive interpretation but were useful in comparing different models or comparing a model’s performance on different classes. We plotted the ROC and PR curves using the PRROC package in R, which interpolated values of precision, recall, true positive rate and false positive rate across a comprehensive range of threshold values.

### Data processing

The basic data processing pipeline entails the creation of spectrograms for each non-overlapping 12-second segment of audio in the dataset and then processing these images with the CNN to provide predictions on which class the image should be assigned (Ruff et al, 2018). During data processing Ruff et al. (2020) used an intermediate step of segmenting the audio into 12-second clips and created spectrograms from those clips. Further testing (Z. Ruff, unpublished data) revealed that an equally effective approach was to generate spectrograms representing 12-second segments directly from the long-form audio, which resulted in far less disk space used. The set of clips representing all non-overlapping 12-second segments takes up as much disk space as the original audio, while the spectrograms representing the same audio uses approximately 2.5% as much disk space. After generating the full set of spectrograms the program processes the images with the pre-trained CNN model and outputs an array of class scores for each one. This array, along with the filename of the image, is written to a text file in comma-separated value format.

For large-scale data processing we used one or more scripts written in Python version 2.7; these scripts carry out the basic steps of the data processing pipeline in sequence. To increase overall speed of data processing we divided the tasks of spectrogram generation and image classification between separate scripts and used additional scripts to divide the original dataset into chunks of roughly equal size to be processed in parallel, which allowed us to take advantage of computers with many processing cores and a large amount of available memory. The basic processing pipeline described here is versatile, but the details of an optimized implementation depend heavily on available computer resources.

### Verification of model output

We used the Ruff et al. (2020) process as guidance to verify output from the CNN, but we made several important modifications. We constructed a list of audio clips to be reviewed based on the class scores assigned to the corresponding images, as contained in the full output file generated through CNN processing. We first extracted all rows with class scores ≥0.25 for northern spotted owl; these are considered potential spotted owl detections first and foremost, regardless of the scores assigned to them for other classes. We then extracted rows with class scores ≥0.95 for any other non-Noise class, which we considered potential detections for the class with the highest score. We extracted these potential detections from the original audio as 12-second clips, which were sorted into directories by target class. Within the directory for a given target class we further divided potential detections by time, creating a folder for each week that an ARU was deployed. We reviewed these short audio clips and corresponding spectrograms and assigned tags for each species detected in each clip. To maximize detections of northern spotted owl, we set a very low detection threshold of 0.25 for this class and reviewed all clips flagged as potential spotted owl detections at this threshold. For all other classes, we needed only verify enough clips (3–5) to confidently establish the presence of the species at a recording station in each week. In most cases this allowed us to quickly establish detection versus non-detection of each target species at each site by week, which feeds directly into the creation of species-specific encounter histories to be used in occupancy analyses.

We found that technicians could comfortably review several thousand clips per day on average, depending on the number of vocal species present and other environmental factors; the level of review effort can be tailored to suit specific research questions as well as the available time and personnel. Shiu et al. (2020) recommend that detection threshold (and consequently recall) be tailored to a realistic level of review effort by examining the relationship between recall and the number of false positives produced per hour of audio. While reviewing CNN output requires some specialized knowledge and skill, we found that technicians could become proficient in the most common sounds in our study areas within 1–2 weeks of supervised work.

### Desktop processing

We created a desktop application to run the CNN on a personal computer through RStudio using the Shiny interface. This application is written in R but makes use of other software, primarily SoX, through system calls. Spectrograms can be generated from audio data and then classified using the pretrained CNN through a straightforward graphical user interface. Required software includes recent versions of Anaconda, R, Rstudio, and Sox. We have not extensively tested the limits of different hardware but have obtained speeds of approximately 100 hours of audio data processed per hour of processing time on inexpensive laptop computers with 2- and 4-core central processing units. See Appendix 1 for detailed instructions on how to install and use the desktop application to process and classify audio data and to extract clips for manual review following processing.

## Results

### Model training

Model performance as reported by the Keras training procedure continued to improve throughout training; we saved the final model configuration after epoch 100 with training loss of 0.0182, validation loss of 0.0139, training accuracy of 0.9954, and validation accuracy of 0.9969. The learning rate stepdown function was invoked a total of 10 times (i.e., as often as was possible given the patience and cooldown periods that we specified), the last being at epoch 97, and the final learning rate was 9.77×10^−7^, a 1,000-fold reduction from the initial learning rate of 0.001. The full training run took approximately 12.5 hours.

### Test set performance

Precision (Fig. 2), recall (Fig. 3) and F1 scores (Fig. 4) for the CNN’s performance on the test set varied for the 16 non-Noise classes across a range of thresholds. Performance was generally stronger for owls than for the other species. Among the owls, precision was highest for northern saw-whet owl and northern pygmy-owl, although precision at higher thresholds exceeded 90% for all owl species except great horned owl, for which precision was noticeably lower at all thresholds (Fig. 2). Among other species, precision was highest for Townsend’s chipmunk, band-tailed pigeon, Steller’s jay, and common raven (Fig. 2). Precision was relatively low for Douglas’ squirrel chirp call, mountain quail, and red-breasted sapsucker (Fig. 2). Precision was lowest for the Douglas’ squirrel rattle call and did not exceed 50% for this class even at the highest threshold of 0.99 (Fig. 2).

**Figure 2.**
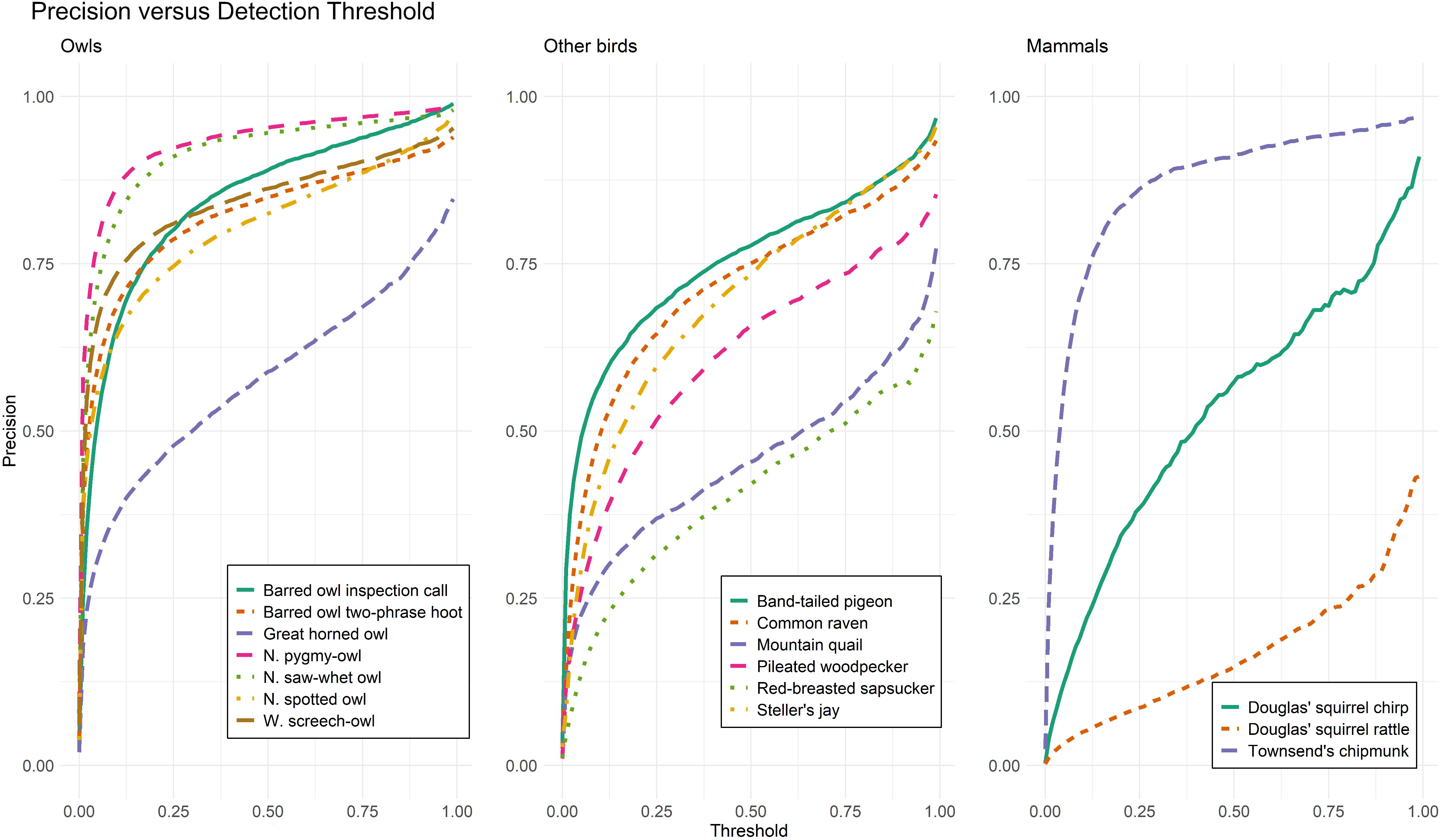
Precision versus detection threshold for 16 sounds produced by 14 avian and mammalian species. Precision or specificity is calculated as [True Positives] / [True Positives + False Positives], considering only clips with class score exceeding the detection threshold for each target class. Precision represents the proportion of apparent “hits” that correspond to real instances of the class in question.

**Figure 3.**
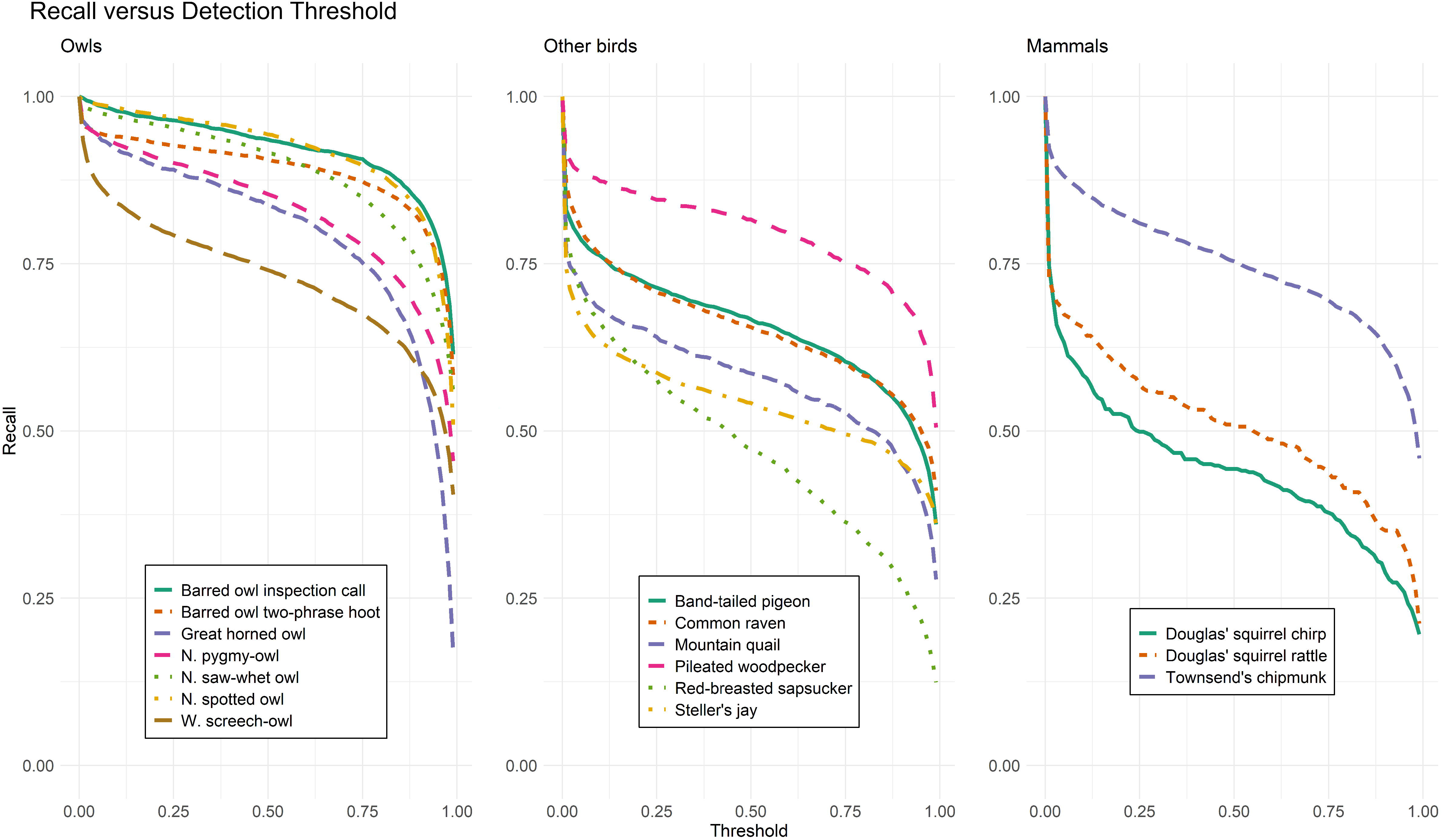
Recall versus detection threshold for 16 sounds produced by 14 avian and mammalian species. Recall or sensitivity is calculated as [True Positives] / [True Positives + False Negatives], considering only clips with class score exceeding the detection threshold for each target class. Recall represents the proportion of real examples present in the dataset that are detected and correctly identified at a given detection threshold.

**Figure 4.**
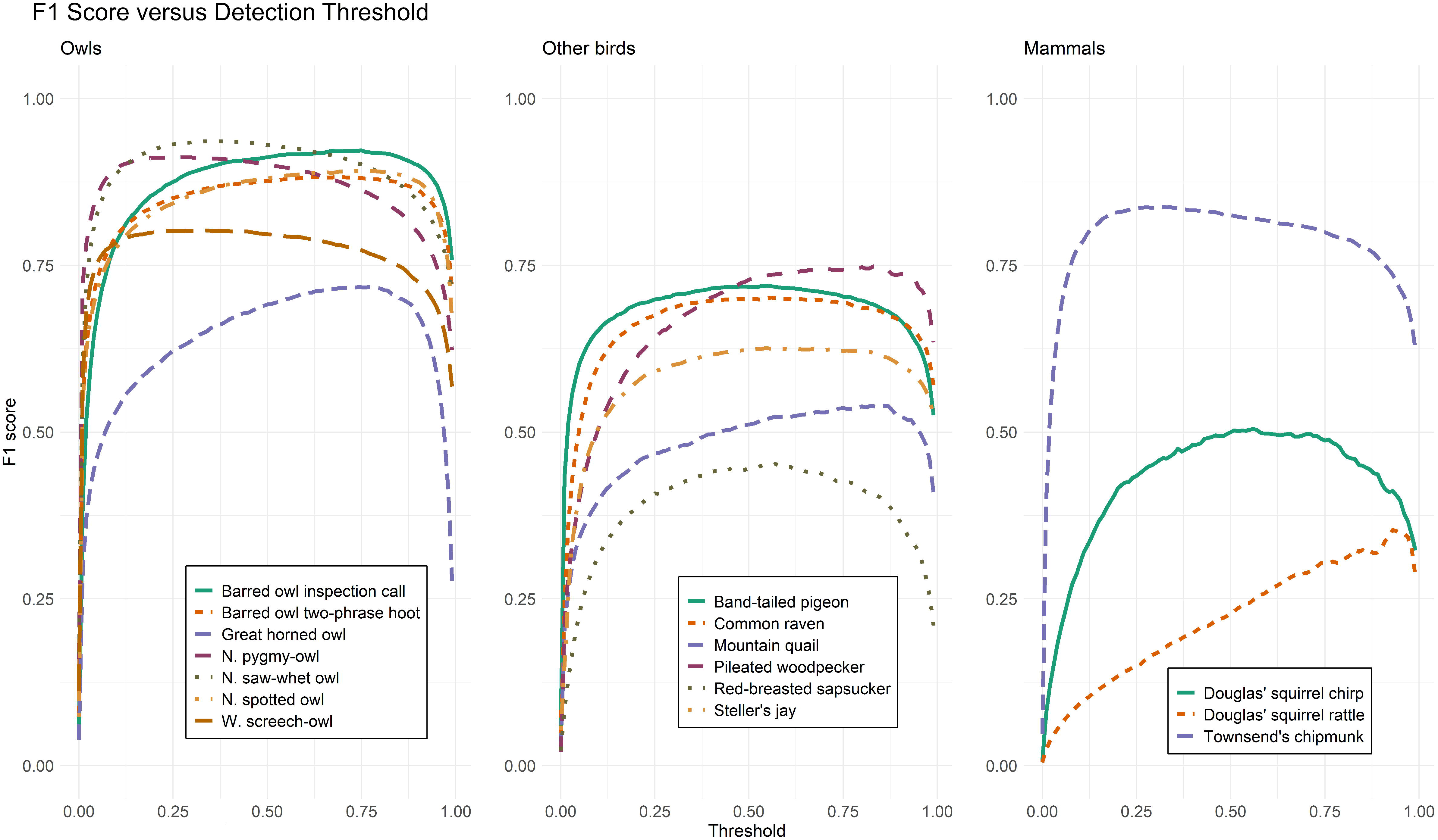
F1 score versus detection threshold for 16 sounds produced by 14 avian and mammalian species. F1 score is calculated as [2 * Precision * Recall] / [Precision + Recall], with both precision and recall calculated at a specific detection threshold. F1 score is intended as a balance of precision and recall and is used to gauge overall model performance.

Among the owl species, recall was best for spotted owl, barred owl (both call types), and northern saw-whet owl, somewhat lower for northern pygmy-owl and great horned owl, and lowest for western screech-owl across most of the range of thresholds, although recall was well above 50% for most species even at thresholds of 0.9 or more (Fig. 3). Recall for other species was less consistent and was highest for pileated woodpecker and Townsend’s chipmunk, moderate for band-tailed pigeon and common raven, lower for mountain quail and Steller’s jay, and lowest for red-breasted sapsucker and Douglas’ squirrel (Fig. 3). Most classes showed recall above 50% at thresholds >0.9.

The plots of F1 score versus threshold indicated that the CNN had a fairly balanced mix of precision and recall for most of the owl classes across a broad range of thresholds, as demonstrated by the flatness of the curves at moderate threshold values (Fig. 4). Great horned owls had markedly better F1 scores at higher thresholds, which may be attributable to the low precision for this class across most of the range of thresholds. F1 score peaks at low threshold values for several species of owl, including northern pygmy-owl, northern saw-whet owl, and western screech-owl (Fig. 4). These species had high precision at low threshold values which appeared to then be offset by diminishing recall at higher thresholds. Similar patterns were visible for non-owl avian species, although these covered a broader range of values. We observed the best F1 scores for pileated woodpecker, common raven, and band-tailed pigeon, depending on threshold (Fig. 4). Mammals showed a wide range of F1 scores; Townsend’s chipmunk was comparable to the owls, while Douglas’ squirrel showed low F1 for the chirp call and lower F1 for the rattle call (Fig. 4).

The ROC curves show close to ideal performance for most owl species, good performance for several of the other bird species including band-tailed pigeon and pileated woodpecker, as well as Townsend’s chipmunk, and somewhat weaker performance for common raven, Steller’s jay, and both Douglas’ squirrel classes (Fig. 5). The AUC values for ROC curves indicated generally good discriminative ability of the CNN across all classes. CNN performance as measured by ROC-AUC was especially high for all owls: barred owl inspection call = 0.998, barred owl two-phrase hoot = 0.973, great horned owl = 0.979, northern pygmy-owl = 0.978, northern saw-whet owl = 0.991, northern spotted owl = 0.995, and western screech-owl = 0.971. ROC-AUC values for most other species also indicated good predictive power; band-tailed pigeon = 0.892, common raven = 0.925, Douglas’ squirrel chirp call = 0.849, Douglas’ squirrel rattle call = 0.838, mountain quail = 0.870, pileated woodpecker = 0.946, red-breasted sapsucker = 0.870, Steller’s jay = 0.818, Townsend’s chipmunk = 0.962.

**Figure 5.**
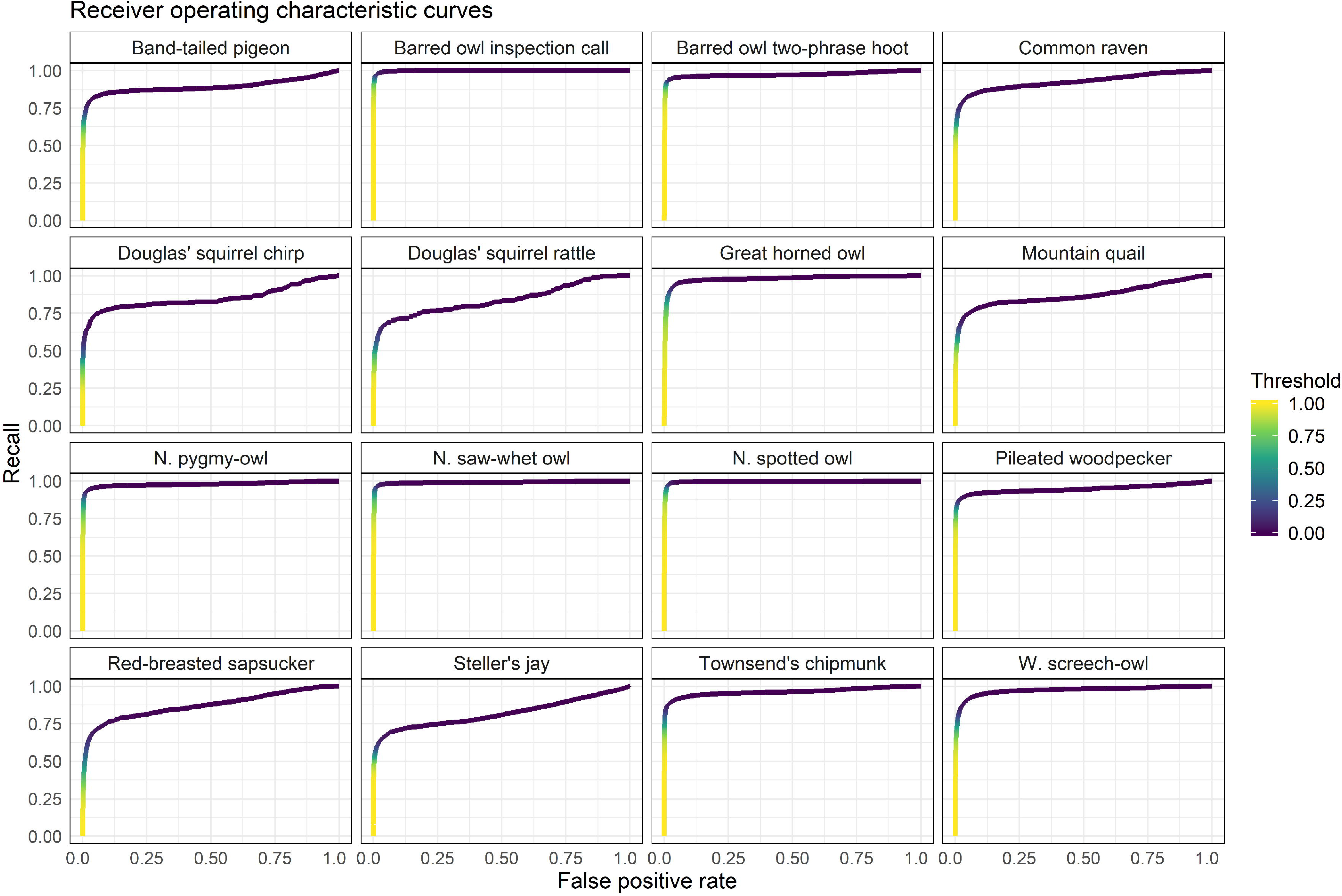
Receiver operating characteristic curves for 16 sounds produced by 14 avian and mammalian species. The receiver operating characteristic curve plots the true positive rate (recall), calculated as [True Positives] / [True Positives + False Negatives] against the false positive rate, calculated as [False Positives] / [False Positives + True Negatives].

The PR curves were more variable than the ROC curves, but showed strong CNN performance for most owls, which was also supported by PR-AUC values: barred owl inspection call = 0.967, barred owl two-phrase hoot = 0.875, great horned owl = 0.715, northern pygmy-owl = 0.945, northern saw-whet owl = 0.961, northern spotted owl = 0.948, and western screech-owl = 0.852. The PR-AUC values for other species showed varied performance: band-tailed pigeon = 0.736, common raven = 0.720, Douglas’ squirrel chirp call = 0.444, Douglas’ squirrel rattle call = 0.267, mountain quail = 0.483, pileated woodpecker = 0.735, red-breasted sapsucker = 0.361, Steller’s jay = 0.602, Townsend’s chipmunk = 0.864 (Fig 6).

**Figure 6.**
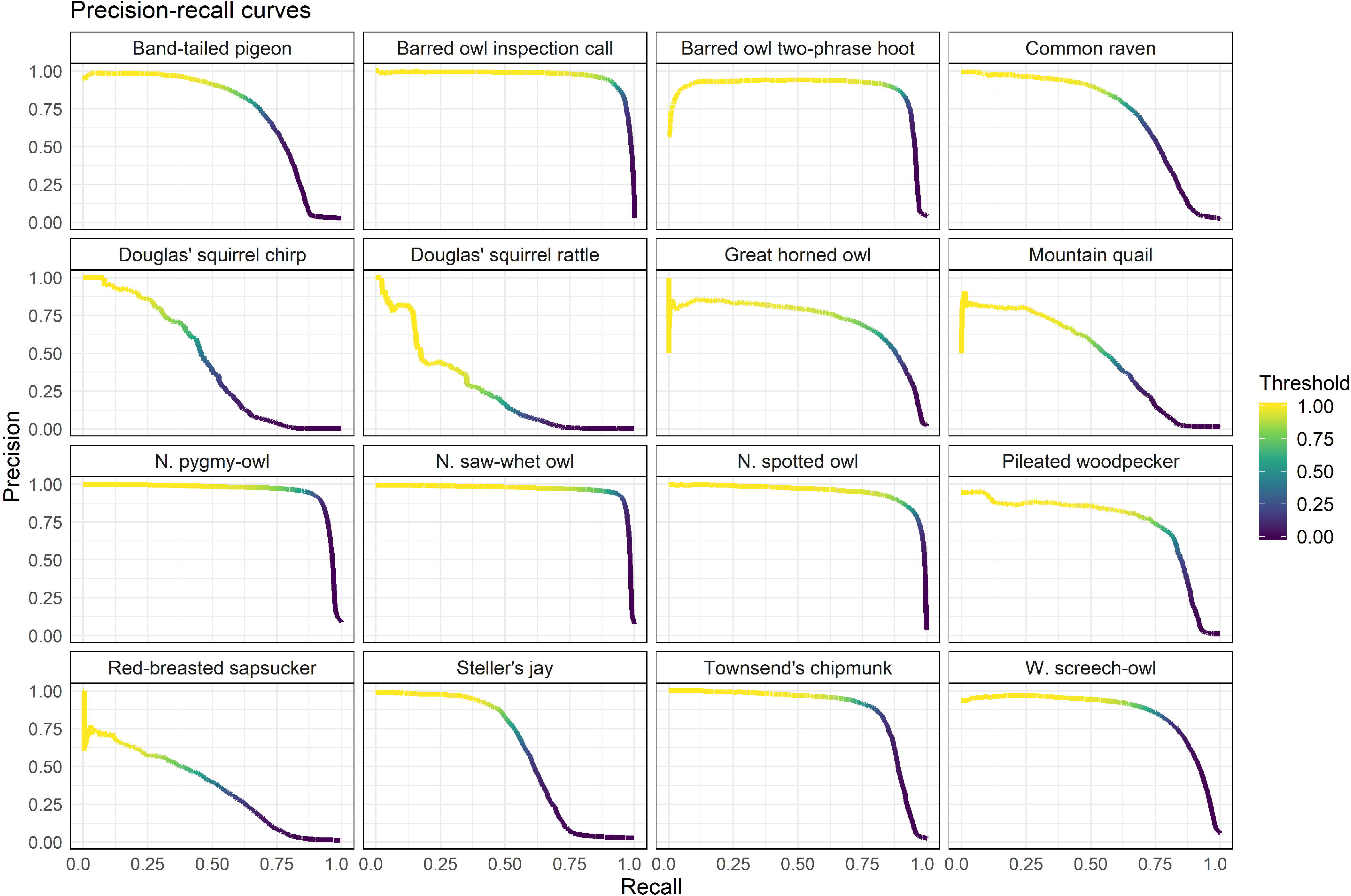
Precision-recall curves for 16 sounds produced by 14 avian and mammalian species. The precision-recall curve plots precision (True positives / [True Positives + False Positives]) against recall (True Positives / [True Positives + False Negatives]) to illustrate a given model’s performance and the magnitude of the trade-off between these metrics across a range of detection thresholds.

## Discussion

Here we present an improvement in CNN performance by retraining with larger set of images and by increasing the number of target classes, resulting in a demonstratively higher-performing CNN compared to the network previously reported in Ruff et al. (2020). Additionally, we packaged the CNN as a desktop application run through Rstudio, making the benefits of this tool available to field biologists and practitioners in a portable and user-friendly interface that requires only free and widely available software. Our CNN had substantially better performance than the Ruff et al. (2020) version for six owl species as demonstrated by area under curve values for both the ROC and the PR curves for each species. To demonstrate the ability to have the CNN distinguish multiple call types for a species we added the barred owl inspection call class, which showed extremely strong performance with distinguishability from the 8-note call, providing progress towards automating detailed biological meaning from passive acoustic data well beyond simple species presence. We also incorporated eight additional species, for which performance was mixed but still strong enough to use CNN output as the foundation for ecological analyses for most target species. Performance for the non-owl classes was comparable to that observed for the original six owl classes as reported by Ruff et al. (2020), which suggests that performance for these classes may improve to a similar degree in future versions. Manual review of apparent detections by humans has the side benefit of producing training data which can be fed back into the network to improve performance in an iterative fashion by periodically retraining the model.

The use of softmax activation in the output layer of the CNN used by Ruff et al. (2020) implied that class labels were exhaustive and mutually exclusive, i.e. each image was implied to have exactly one correct label. This is not strictly true because in natural systems it is not unusual for multiple species to vocalize simultaneously. Because the number of target species for the Ruff et al. (2020) CNN was relatively small, the inability to recognize multiple classes in the same image was unlikely to be a major limitation, since single-class images would likely outnumber multi-class images in most cases. However, the lists of target species are likely to expand with future CNN developments, so multiple target classes occurring in some images will be more common. As such it will be increasingly important that these multi-class events be accurately captured in the model output as multi-class predictions. The use of sigmoid activation in the output layer of the reported CNN should allow for multi-label classification, in which a single image can receive high scores for multiple classes. In practice, we did not find that the CNN reliably assigned high scores to each appropriate class when multiple species were present in an image. This may be because our training set contained only singly labeled images. These results suggest that effort should be made to include images with multiple correct labels – perhaps even generating them artificially by combining multiple single-class images – to train the CNN to more reliably recognize multi-class images. Alternatively, recent work has obtained strong multi-label performance using “pseudo-labeled” training data, in which each training example has one class labeled as present or absent and all other classes labeled as unknown, and a custom loss function which penalizes incorrect prediction for only the labeled class (Zhong et al. 2020).

We found that recall for non-owl birds and mammals was well below 100% even at very low thresholds (e.g. 0.05), suggesting that the CNN assigned a very low score to the correct class in a significant number of these cases. This was also true for western screech-owl, though less dramatically so. A larger proportion of test images of the non-owl classes featured calls of multiple species and therefore had multiple correct labels. Because the CNN did not reliably assign high scores to every class that was present in multi-class images, the lower recall observed for non-owls may be an artifact of our relatively modest test set rather than a feature of the CNN itself.

Precision was noticeably weak for the Douglas’ squirrel rattle call even at high thresholds. This may be attributable to both the character of the call itself and our specific processing pipeline. The rattle call consists of an extended sequence of rapidly repeated (~15 s^−1^) chirps. The speed of the call combined with the time resolution of our spectrogram images (500 pixels representing 12 s of recording time) means that even in cases with a high signal-to-noise ratio, individual chirps may be separated by as little as one pixel, and this separation may effectively vanish when combined with echoes, scattering, and ambient sounds. This call is also highly variable in length, although this does not seem to have hindered classification for other calls such as Townsend’s chipmunk or northern saw-whet owls. It is possible that our training set for this species simply did not establish a sufficiently distinctive pattern, given the resolution of the images, to enable the CNN to disregard similar sounds. Common sources of false positives for the Douglas’ squirrel rattle call were wind, insects, and anuran calls. In spite of some noted issues, the performance of our CNN was broadly comparable to that of another recent CNN which achieved precision ranging from 0.13 to 1.00 and recall ranging from 0.25 to 1.00 at a detection threshold of 0.99 for 24 classes of avian and anuran vocalizations from Puerto Rico (LeBien et al. 2020).

With our development of the Shiny App this model is quite portable and can be run by biologists with relatively short turnaround time for data processing. While training deep convolutional neural networks is computationally intensive and benefits from high-performance computer hardware, particularly powerful graphics processing units, the actual processing of audio data can be done at a reasonable speed on consumer-grade computers. Because the task of generating spectrograms can be parallelized to a substantial degree, this task makes efficient use of multi-core processors, and the availability of inexpensive 8- and 12-core central processing units makes desktop processing increasingly attractive. Classifying the resulting images is not computationally demanding and can be run at reasonable speed without relying on powerful graphics processing units. Depending on the specific hardware configuration, processing speed may be limited either by the data connection or the read-write speeds of the storage media. We have obtained the best processing speeds with data stored on internal solid-state drives with high-speed data connections; however, external hard drives connected by universal serial bus still offers satisfactory performance. The desktop application also included functions for extracting rows representing potential detections from the raw results file and for extracting short clips corresponding to these rows for subsequent verification.

Moving beyond the ability to process audio on consumer-grade computers, the advantages of large-scale passive acoustic monitoring may only be fully realized when the raw data can be processed in close to real time (i.e., as they are collected) and salient results communicated quickly to biologists and managers. This may entail processing data in a distributed fashion using small, inexpensive system-on-chip processing devices coupled to the recording device, or the use of purpose-made recording devices with software for onboard processing, e.g. AudioMoth (Hill et al. 2017, Prince et al. 2019). Such distributed processing nodes could communicate potential detections to biologists remotely over mobile data networks, streamlining the process of retrieving data from the field and allowing for very rapid responses to emergent issues at field sites (e.g. the Rainforest Connection project; https://rfcx.org). However, engineering such an all-in-one solution to remain active for weeks at a time and to withstand the environmental conditions typical of many field sites is a non-trivial challenge. These developments will require multi-disciplinary collaborations between ecologists, computer scientists, and engineers. From a species conservation and management standpoint these advancements will be crucial to enhance our ability to monitor target species in close to real time. This is especially true for species such as northern spotted owls which are rare, elusive, and have habitats that are often subject to land management actions with economic and ecological implications.

## Acknowledgments

Funding for this research was provided by the USDA Forest Service and USDI Bureau of Land Management, and we thank R. Davis, B. Hollen, and G. McFadden for facilitating that funding. We thank the many field biologists and lab technicians that collected and validated much of the data presented here, including C. Cardillo, D. Culp, L. Duchac, Z. Farrand, T. Garrido, E. Guzman, A. Ingrassia, D. Jackobsma, E. Johnston, R. Justice, K. McLaughlin, A. Munes, P. Papajcik, J. Runjaic, and S. Sabin. The primary computer system used for the development of the convolutional neural network was owned and administered by the Center for Genome Research and Biocomputing at Oregon State University. The findings and conclusions in this publication are those of the authors and should not be construed to represent any official U.S. Department of Agriculture or U.S. Government determination or policy. The use of trade or firm names in this publication is for reader information and does not imply endorsement by the U.S. Government of any product or service.

## Notes

### Competing Interest Statement

The authors have declared no competing interest.

