## Appendix 1 for "A Convolutional Neural Network and R-Shiny App for Automated Identification and Classification of Animal Sounds"

**Installation and use of CNN for desktop processing with Shiny and Rstudio**

### Installing required software

For purposes of this guide we assume that you are using a computer running a 64-bit version of Windows on which you can install programs, i.e. you will need either an account with administrator privileges or at least the ability to run programs as an administrator or elevated. Keras will not function on 32-bit operating systems.

**Step 1.** Install the most recent version of R. If it is already installed, go to step 2.

Go to https://cran.rstudio.com/ and click "Download R 3.x.x for Windows". (Latest version at time of writing was R 3.6.2.) Open your Downloads folder and run the installer. It will be called something like R-3.6.2-win.exe. All the default settings should be fine.

**Step 2.** Install Rstudio. If it is already installed, go to step 3.

Go to https://rstudio.com/products/rstudio/download/#download and click the big blue button that says "Download Rstudio for Windows". This file is about 150 MB. Go to your Downloads folder and run the installer. Again, the default settings will work.

**Step 3.** Update R and Rstudio.

If you just installed both programs, skip this step. Updating regularly is good practice.

To update R, open the R console. Run the following commands:

```
install.packages("installr")
library(installr)
updateR()
```

This will install the "installr" package, load the package, and run the updateR function, which will search for the latest version and install it if there is one available. It will generate various popups; just agree to the default options and it should all be fine.

To update Rstudio, open Rstudio. In the menu bar up top, click Help > Check for Updates. If there is a newer version available, click "Quit and download". This may not be strictly necessary as long as your packages are reasonably up to date, but if you have been using the same version for a year or more, you should probably update regardless.

**Step 4.** Install SoX.

Sox is a small command-line program that we use to process audio data. You won't need to do anything directly with it, but it needs to be installed on your computer. The specific version doesn't matter too much. Go to https://sourceforge.net/projects/sox/files/sox/ and click "Download latest version" (the big green button) to download the executable installer for Windows. Open your Downloads folder and run the installer. Make a note of where it is installed; the default is something like C:\Program Files (x86)\sox-14-4-2.

**Step 5.** Install Anaconda for Windows.

Anaconda is a massive set of software tools intended for scientific use. Go to https://www.anaconda.com/distribution/ and scroll down to where it says "Anaconda xxxx.xx for Windows Installer." Under "Python 3.x version" click the green button that says Download. This will download the installer for the 64-bit Windows version. The download is a few hundred MB so it may take a few minutes.

**Step 6.** Install Keras for R.

Keras is an application programming interface (API) that allows R to talk to the TensorFlow software library to build and run neural networks. Open Rstudio. In the Packages tab in the lower-right-hand pane, click Install. In the pop-up window, under Packages, type "keras" and click Install. You may get a warning in the console but as long as there are no error messages it should be fine. Next, in the console, type `library(keras)` and hit Enter. R will load the Keras library. Now, type `install_keras()` and hit Enter. This will tell R to configure keras and all its dependencies. There will be a lot of output in the console window and most of it will mean nothing to you. Again, as long as there are no error messages, it should work.

**Step 7.** Install the shiny, tidyverse, and tuneR packages.

These packages define the functioning of the graphical user interface to the CNN app and provide some helpful functions, particularly with respect to the handling of WAV files. Rstudio should still be open. Again, in the lower-right corner, under Packages, click Install, and enter the names of each package. Notice that you can tell R to install more than one at a time if you separate the package names with commas or spaces. Again, there will be a lot of output in the console window, which you shouldn't need to worry about.

**Step 8.** Close Rstudio and open it again.

At this point the CNN application should be usable and you should have everything you need to process audio data. See below for instructions for using the app.

**Step 9.** (Recommended) Install Bulk Rename Utility.

As the name implies, this is a simple utility for quickly and efficiently renaming many files. The app assumes that files are named according to the same convention, which encodes some important information in the filename. It is important that they be formatted correctly.

### **Filename formatting**

The format your audio files should be under is as follows:

Area_Site-Station_Date_Time.wav

This filename consists of four components separated with underscores: Study area, specific location, date, and time. The Study area or project ID is typically a three-letter code. Site and station ID should both be alphanumeric and should be separated by a dash (-). The date should be in YYYYMMDD format. The time should be in HHMMSS format in 24-hour time. The time and date should already be in the correct format if the audio is from Song Meter ARUs.

For instance, our filenames are formatted as follows:

COA_8908-5_20190716_201530.wav

The filename indicates that this file was recorded in the Oregon Coast Range study area (COA), in hexagon 8908, at station 5, and was recorded beginning on July 16, 2019, at 8:15:30 PM. Note that both the spatial and temporal information in this filename is arranged in order from most general to most specific – from study area to hexagon to station within the hexagon, and from year to month to day, and finally from hour to minute to second.

### Using the CNN through the Shiny interface

These instructions assume you have installed all the necessary software as outlined above.

Note: This application works for our purposes, and we have tried to make it somewhat user-friendly, but it was not made by professional software engineers, so its operation is somewhat finicky. It makes some assumptions in order to work efficiently, and it may not work well (or at all) if those assumptions are broken. Be careful!

You will launch this application from within Rstudio by typing a commands into Console pane; further operation will use the Shiny window that appears once the program is running. The Shiny window will take over Rstudio's input and output as long as it is open; you will not be returned to the normal prompt until this window is closed. Some of the program's output (progress bars, messages, etc.) will appear in the console pane, so it is helpful to keep an eye on Rstudio while the CNN application is running.

**Step 1.** Download the "Shiny_CNN" folder from the Zenodo page at https://doi.org/10.5281/zenodo.3923030 and copy the folder to your normal home directory in Rstudio, i.e. whatever folder comes up if you open Rstudio and run `getwd()`. This home directory needs to contain a folder called "Shiny_CNN" which should contain an R file called app.r, a text file called "readme" which lays out some potentially helpful information, and another file called Owl_CNN.h5, which is the pretrained CNN model.

**Step 2.** Open Rstudio. In the console window, type `shiny::runApp("Shiny_CNN")` and hit Enter. Rstudio will search the current working directory for a folder called "Shiny_CNN" containing a file called app.r; if it does not find these items, you will get an error. Asssuming the folder and app file are found, a Shiny application window will appear. You will also get some output in the console pane as the app loads some R packages, followed by the message "Listening on http://127.0.0.1:xxxx", which indicates that Rstudio is communicating with the app through a socket on the local machine. When first launched, the Shiny window will look like this:

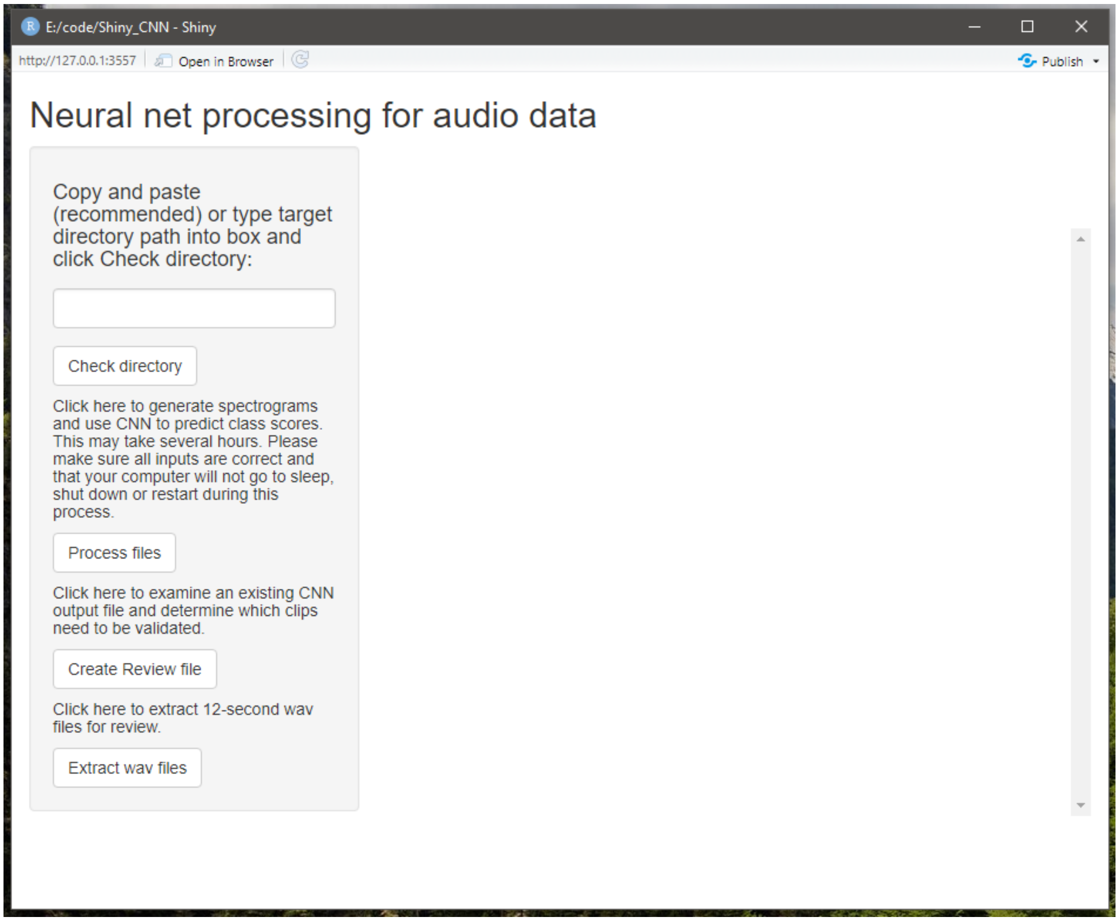


**Step 3.** In the Shiny window, use the text box on the left-hand side to input the directory containing the files that you want to process. Use the "Check directory" button to verify that R can find some WAV files. The number that shows up is wav files that are ≥500 KB in size and do not have "_part_" in the name. You will get a list of the files that have been found. The app will also tell you if there is already a CNN results file and/or a Review file in the directory.

**Step 4.** Click the button labeled Process files. Be careful to only click this button once! If you give the program input while it is busy, the input will be stored and then executed when the current task finishes. Clicking the button multiple times may cause your data to be processed multiple times in sequence, which can waste a great deal of time.

Note that you may also get a Windows Firewall pop-up saying that features of the program have been blocked. You can close this pop-up; the program will continue to run.

This will first create spectrograms in the form of image files in Portable Network Graphic (.png) format, representing non-overlapping 12-second segments of the audio. The images will be created in a new directory called Temp/images within the folder you specified in step 4. There will be 300 images for every hour of audio, so if you have a substantial dataset there will be a lot of them. This will likely be the most time-consuming part of the process; we recommend using a computer with multiple cores. We have not implemented a progress bar for this step, but as a crude workaround you can open the Temp/images folder in Windows Explorer and watch as the files are created.

Note that this step is by far the longest; processing time will vary depending on your computer's hardware capabilities and where the data are stored. As a *very* general rule of thumb, with data stored on external USB hard drives, you should be able to process approximately 100 hours of recordings per hour of processing time.

Once all the images have been generated, the app will load the trained CNN model and start classifying the images in batches. Keras will create a text-based progress bar in the console pane in Rstudio showing how many batches have been processed. Once the full set of images has been classified the app will write the results to a file in the same folder. Once you get a message telling you that the output file has been written (`"Output written to CNN_Predictions_[location].csv in folder D:/Path/To/Folder"`) you can assume that this step is done and the program is ready for more input. As the results file is in comma-separated value (.csv) format, it can be opened in Microsoft Excel or your favorite text editor. The Shiny window will not immediately display the results file; you will have to click Check directory again so the app can find it.

**Step 5.** Click the button labeled Create Review file. This will search the CNN results file for lines where a) the score for spotted owl (STOC) was ≥0.25 or b) the score for any of the other target classes besides Noise was ≥0.95 and write these lines to a new file, along with a randomly chosen subset of 0.5% of the other lines. This is the set of clips that will be extracted for human review under our current validation procedure. If all goes well the app will write a message to the console pane that says `Creating review file based on D:/Path/To/CNN_Predictions_[location].csv…` followed by `Done` when it finishes. This step is very quick. Again, the review file will not appear in the Shiny window until you click Check directory.

**Step 6.** To extract the clips listed in the review file, click Extract wav files. This will create a new folder called Review in the folder specified in step 4. The Review folder will have subfolders for each of the target classes, which will contain the clips of that class that warrant further review.

Depending on the dataset and how much other data is on the same hard drive, disk space may be an issue when extracting clips for review. Each 12-second clip is about 1.5 MB, so every 10,000 such clips will take up about 15 GB. You can use the number of lines in the Review file to judge how much free space you will need before extracting clips for review.

The app will print a message to the console that says "`Extracting wav files beginning at [time]…`" Extracting the clips may take some time. When all the new clips have been made there will be another message that says "`Finished at [time]. [n] clips extracted.`" Once you get the message that review clips have been extracted, you can close the Shiny window and exit Rstudio. Below is an example of console output from the processing of a very small test set.

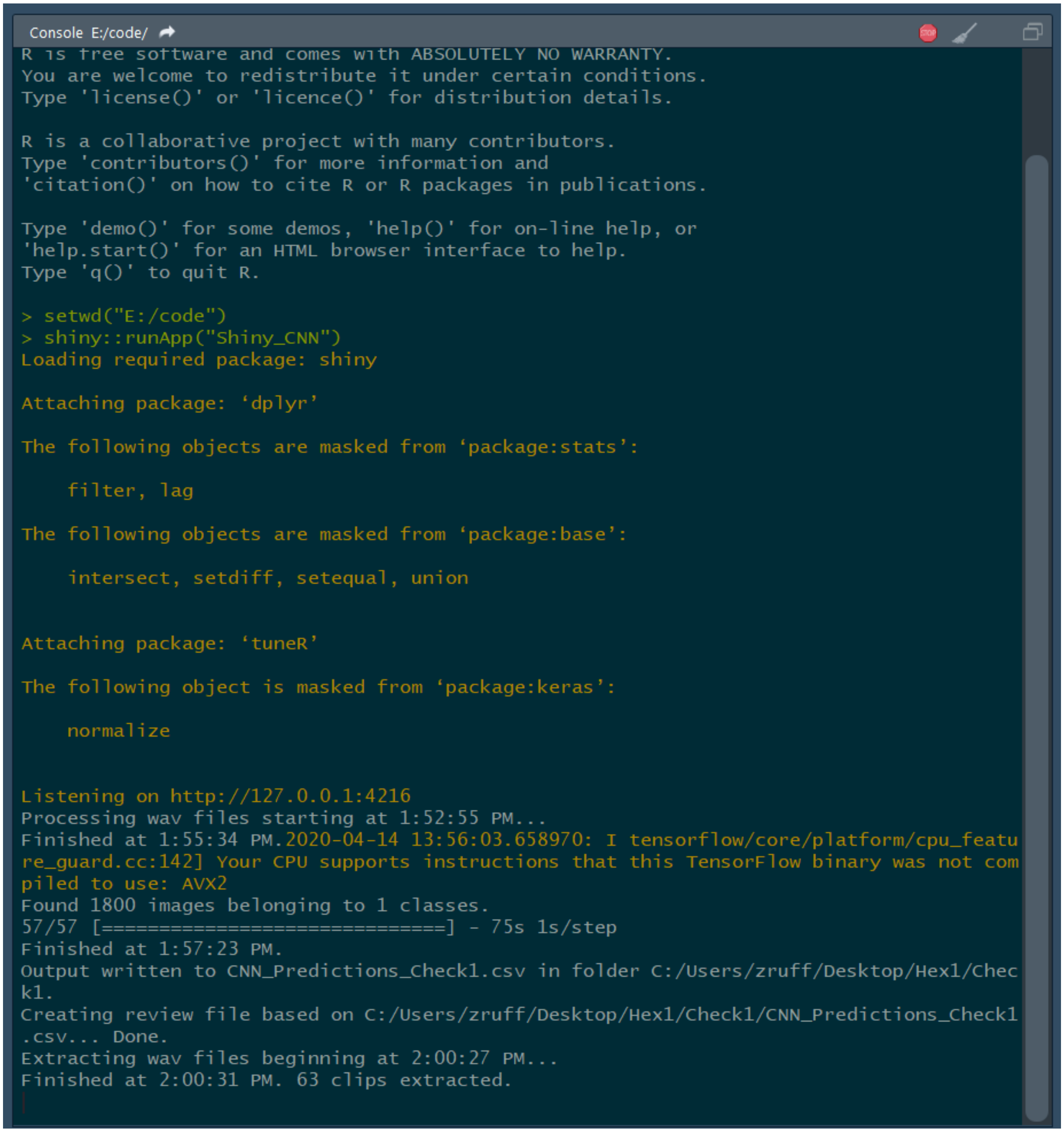

```
Console E:/code/
R is free software and comes with ABSOLUTELY NO WARRANTY.
You are welcome to redistribute it under certain conditions.
Type 'license()' or 'licence()' for distribution details.

R is a collaborative project with many contributors.
Type 'contributors()' for more information and
'citation()' on how to cite R or R packages in publications.

Type 'demo()' for some demos, 'help()' for on-line help, or
'help.start()' for an HTML browser interface to help.
Type 'q()' to quit R.

> setwd("E:/code")
> shiny::runApp("Shiny_CNN")
Loading required package: shiny

Attaching package: ‘dplyr’

The following objects are masked from ‘package:stats’:

    filter, lag

The following objects are masked from ‘package:base’:

    intersect, setdiff, setequal, union


Attaching package: ‘tuneR’

The following object is masked from ‘package:keras’:

    normalize


Listening on http://127.0.0.1:4216
Processing wav files starting at 1:52:55 PM...
Finished at 1:55:34 PM.2020-04-14 13:56:03.658970: I tensorflow/core/platform/cpu_featu
re_guard.cc:142] Your CPU supports instructions that this TensorFlow binary was not com
piled to use: AVX2
Found 1800 images belonging to 1 classes.
57/57 [==============================] - 75s 1s/step
Finished at 1:57:23 PM.
Output written to CNN_Predictions_Check1.csv in folder C:/Users/zruff/Desktop/Hex1/Chec
k1.
Creating review file based on C:/Users/zruff/Desktop/Hex1/Check1/CNN_Predictions_Check1
.csv... Done.
Extracting wav files beginning at 2:00:27 PM...
Finished at 2:00:31 PM. 63 clips extracted.
```

Once clips for review have been extracted, you can close the Shiny window by clicking the X in the upper right-hand corner or by pressing Esc from the Rstudio console window. The actual review of these clips can be done with various programs but is outside the scope of this paper.
